# UMAP does not preserve global structure any better than t-SNE when using the same initialization

**DOI:** 10.1101/2019.12.19.877522

**Authors:** Dmitry Kobak, George C. Linderman

## Abstract

One of the most ubiquitous analysis tools employed in single-cell transcriptomics and cytometry is t-distributed stochastic neighbor embedding (t-SNE) [1], used to visualize individual cells as points on a 2D scatter plot such that similar cells are positioned close together. Recently, a related algorithm, called uniform manifold approximation and projection (UMAP) [2] has attracted substantial attention in the single-cell community. In *Nature Biotechnology*, Becht et al. [3] argued that UMAP is preferable to t-SNE because it better preserves the global structure of the data and is more consistent across runs. Here we show that this alleged superiority of UMAP can be entirely attributed to different choices of initialization in the implementations used by Becht et al.: t-SNE implementations by default used random initialization, while the UMAP implementation used a technique called Laplacian eigenmaps [4] to initialize the embedding. We show that UMAP with random initialization preserves global structure as poorly as t-SNE with random initialization, while t-SNE with informative initialization performs as well as UMAP with informative initialization. Hence, contrary to the claims of Becht et al., their experiments do not demonstrate any advantage of the UMAP algorithm *per se*, but rather warn against using random initialization.

---

At the core of both t-SNE and UMAP are loss functions which make similar points attract each other and push dissimilar points away from each other. Both algorithms minimize their loss functions using gradient descent. Gradient descent begins with some initial configuration of points, and with each iteration, the points are moved to decrease the loss function. The specific implementations of these algorithms used by Becht et al. differed in how the initial configuration of points was chosen: t-SNE implementations placed the points randomly, while the UMAP implementation used Laplacian eigenmaps (LE) [4], an algorithm that can often achieve a good embedding on its own. The effect of this difference on how well the two algorithms preserve global structure was not discussed or investigated by Becht et al.

Using the code published by Becht et al., we analyzed the separate effects of initialization and algorithm by adding UMAP with random initialization and t-SNE (using FIt-SNE [5]) with principal component analysis (PCA) initialization [6] to the benchmarking comparison. For t-SNE, we used PCA instead of LE mainly for computational simplicity (we scaled the PCA initialization to have variance 0.0001 which is the default variance of random initialization in t-SNE [6]). Apart from the initialization, both algorithms were run with default parameters, as in Becht et al. We used all three datasets analyzed in the original publication (sample sizes from 320 000 to 820 000 cells) [7,8,9]. To quantify preservation of global structure, Becht et al. computed Pearson correlation between pairwise distances in the highdimensional space and in the embedding. To quantify reproducibility of the embedding, they embedded random subsamples of the data and measured the correlation of coordinates of subsample embeddings with the coordinates of the full dataset embeddings (up to symmetries across the axes). Our results show that t-SNE and UMAP with random initializations perform similarly poorly with regard to both metrics, while t-SNE and UMAP with LE/PCA initializations perform similarly well (Table 1). See Extended Data Figures 1–5 for the exact analogues of the original figures from Becht et al.

**Table 1:** Performance of t-SNE and UMAP with random and informative initializations, using data sets and evaluation metrics from Becht et al. For the reproducibility metric, the average over three random subsamples of size *n*= 200000 is reported.

|  | Preservation of pairwise distances |  |  | Reproducibility of large-scale structures |  |  |
| --- | --- | --- | --- | --- | --- | --- |
|  | Samusik | Wong | Han | Samusik | Wong | Han |
| UMAP, LE init. | **0.70** | 0.58 | **0.31** | 0.94 | **0.98** | 0.49 |
| UMAP, random init. | 0.40 | 0.39 | 0.14 | 0.24 | 0.21 | 0.22 |
| t-SNE, PCA init. | 0.60 | **0.66** | 0.29 | **0.95** | **0.98** | **0.92** |
| t-SNE, random init. | 0.32 | 0.37 | 0.18 | 0.29 | 0.33 | 0.06 |

Becht et al. wrote that their findings were “consistent with the idea” that UMAP performs “optimizations that are sensitive to global features of the data, thus reaching similar arrangements more consistently”. Our results show that this conclusion is misleading: their findings were in fact not due to UMAP *optimizations* but due to its *initialization.*

We have recently argued that t-SNE with PCA initialization produces more meaningful embeddings than t-SNE with random initialization [6]. The findings of Becht et al. also underscore the importance of using informative initializations, and suggest that it should be used as the default option in t-SNE and UMAP implementations (as is already the case e.g. in `openTSNE` [10], a Python re-implementation of FIt-SNE). Importantly, t-SNE with non-random initialization should not be considered a new algorithm or even an extension of the original t-SNE; it is exactly the same algorithm with the same loss function, and almost any existing implementation trivially allows to use any given initialization, including the PCA-based one.

Our aim here is not to argue which algorithm, t-SNE or UMAP, is more suitable for single-cell studies. Once informative initializations are used, both algorithms seem to preserve the global structure similarly well, and modern implementations of both algorithms work similarly fast (the widespread opinion that UMAP is much faster than t-SNE is outdated: for 2D embeddings, FIt-SNE works at least as fast [3,6]). When comparing the resulting embeddings (Extended Data Figure 6), the most striking difference is that UMAP produces denser, more compact clusters than t-SNE, with more white space in between. Very similar embeddings can be produced by t-SNE with so called exaggeration that increases the attractive forces by a constant factor [6]. Future research in machine learning is needed to pinpoint the exact mathematical and algorithmic origins of this difference, while future research in single-cell biology is needed to decide which algorithm is more faithful to the underlying biological data.

## Author contributions

The authors contributed equally.

## Acknowledgments

The authors thank Philipp Berens, Stefan Steinerberger, and Yuval Kluger for discussions and helpful comments. DK was supported by the Deutsche Forschungsgemeinschaft (BE5601/4-1 and the Cluster of Excellence “Machine Learning — New Perspectives for Science”, EXC 2064, project number 390727645), the Federal Ministry of Education and Research (FKZ 01GQ1601 and 01IS18039A) and the National Institute of Mental Health of the National Institutes of Health under Award Number U19MH114830. GCL was supported by the National Human Genome Research Institute (F30HG010102) and U.S. NIH MSTP Training Grant T32GM007205. The content is solely the responsibility of the authors and does not necessarily represent the official views of the National Institutes of Health.

## Competing interests

The authors declare no competing interests.

## Code availability

The code will be made available at https://github.com/linqiaozhi/DR_benchmark_initialization.

## Extended Data Figures

**Extended Data Figure 1:**
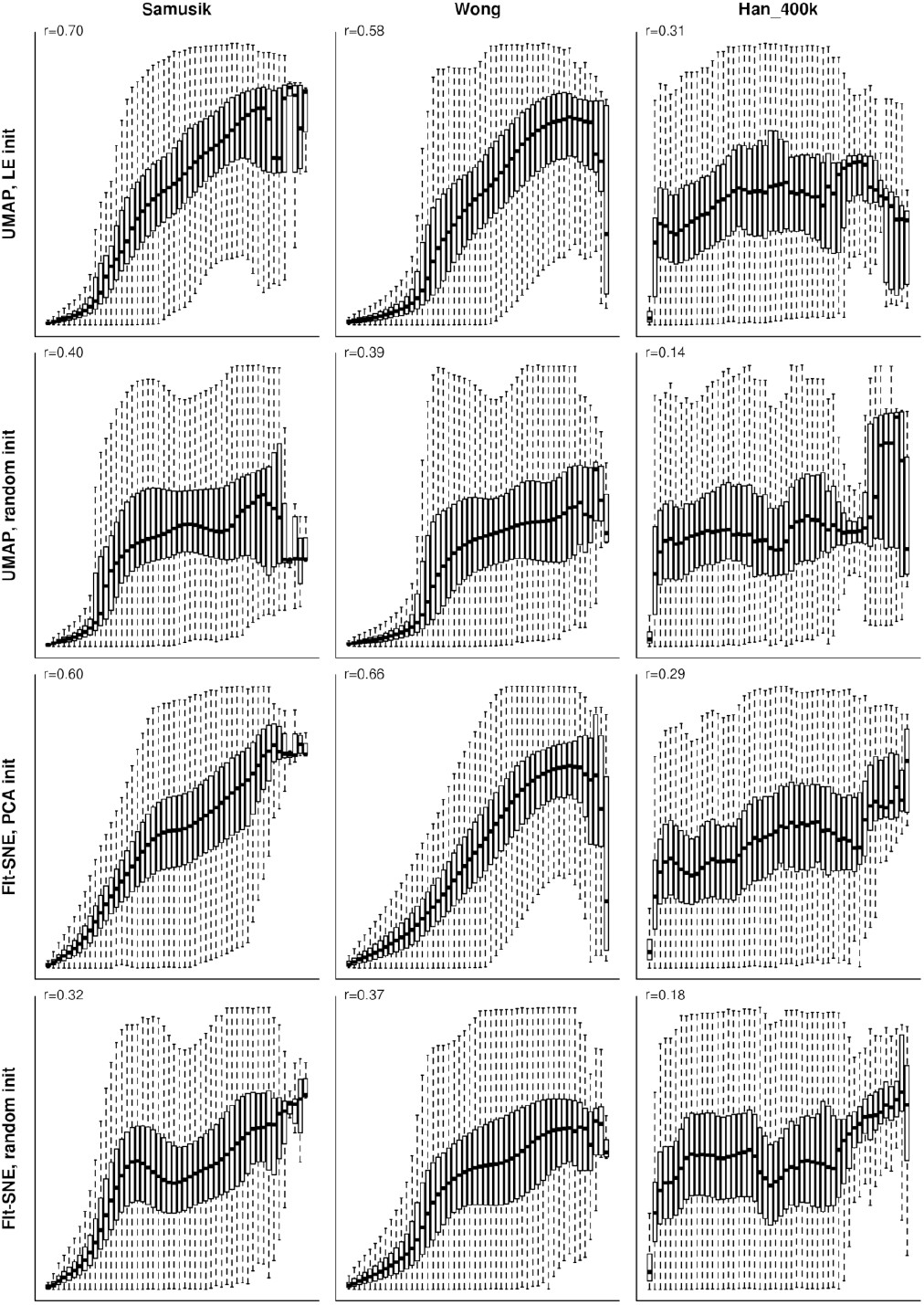
Preservation of pairwise distances in embeddings. The exact analogue of Figure 5 in the original publication. To quote the original caption: “Box plots represent distances across pairs of points in the embeddings, binned using 50 equal-width bins over the pairwise distances in the original space using 10,000 randomly selected points, leading to 49,995,000 pairs of pairwise distances. […] The value of the Pearson correlation coefficient computed over the pairs of pairwise distances is reported. For the box plots, the central bar represents the median, and the top and bottom boundary of the boxes represent the 75th and 25th percentiles, respectively. The whiskers represent 1.5 times the interquartile range above (or, respectively, below) the top (or, respectively, bottom) box boundary, truncated to the data range if applicable.”

**Extended Data Figure 2:**
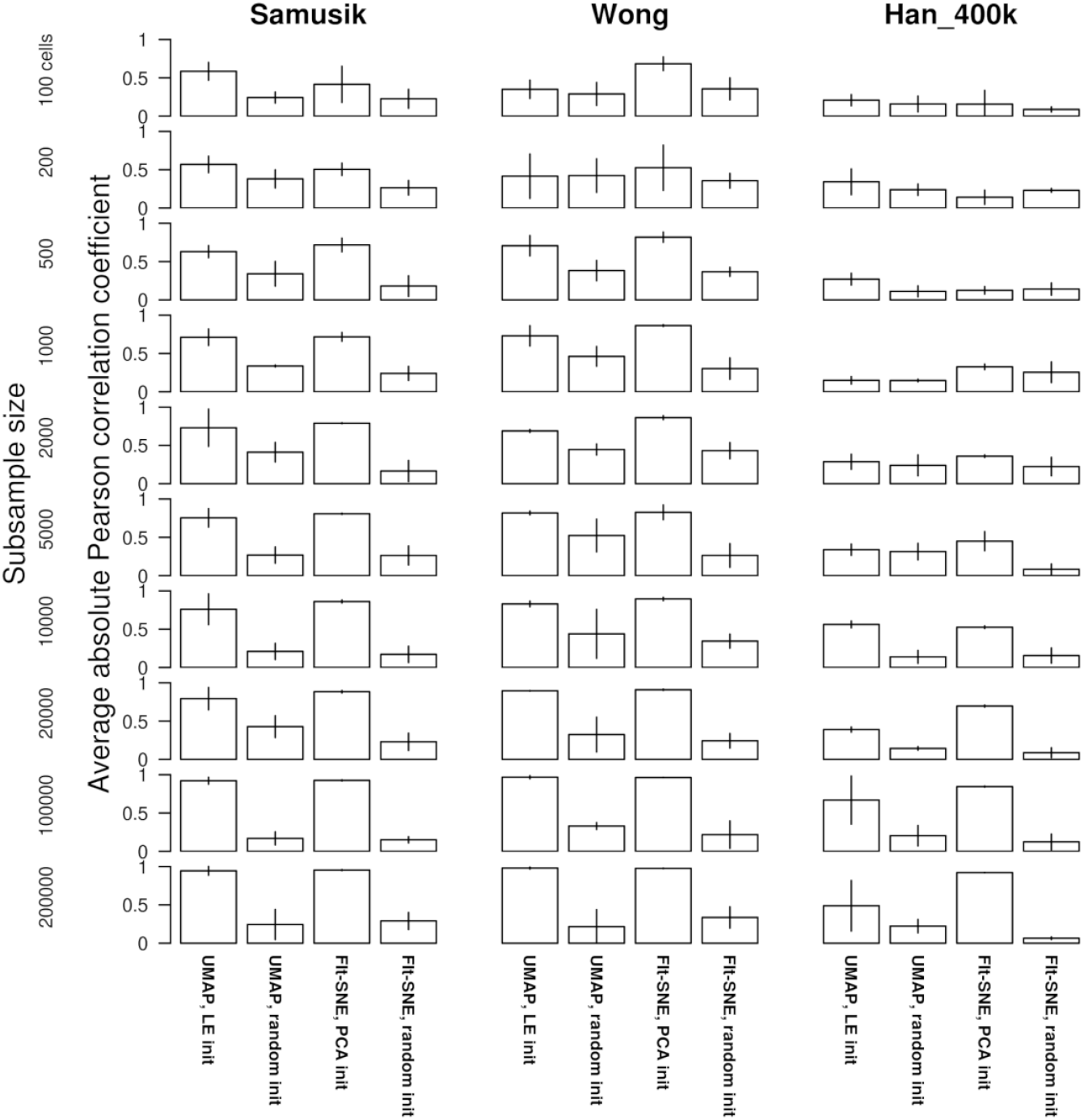
Reproducibility of large-scale structures in embeddings. The exact analogue of Figure 6 in the original publication. To quote the original caption: “Bar plots represent the average unsigned Pearson correlation coefficient of the points’ coordinates in the embedding of subsamples versus in the embedding of the full dataset, thus measuring the correlation of coordinates in subsamples versus in the embedding of the full dataset, up to symmetries along the graph axes. Bar heights represent the average across three replicates and vertical bars the corresponding s.d.”

**Extended Data Figure 3:**
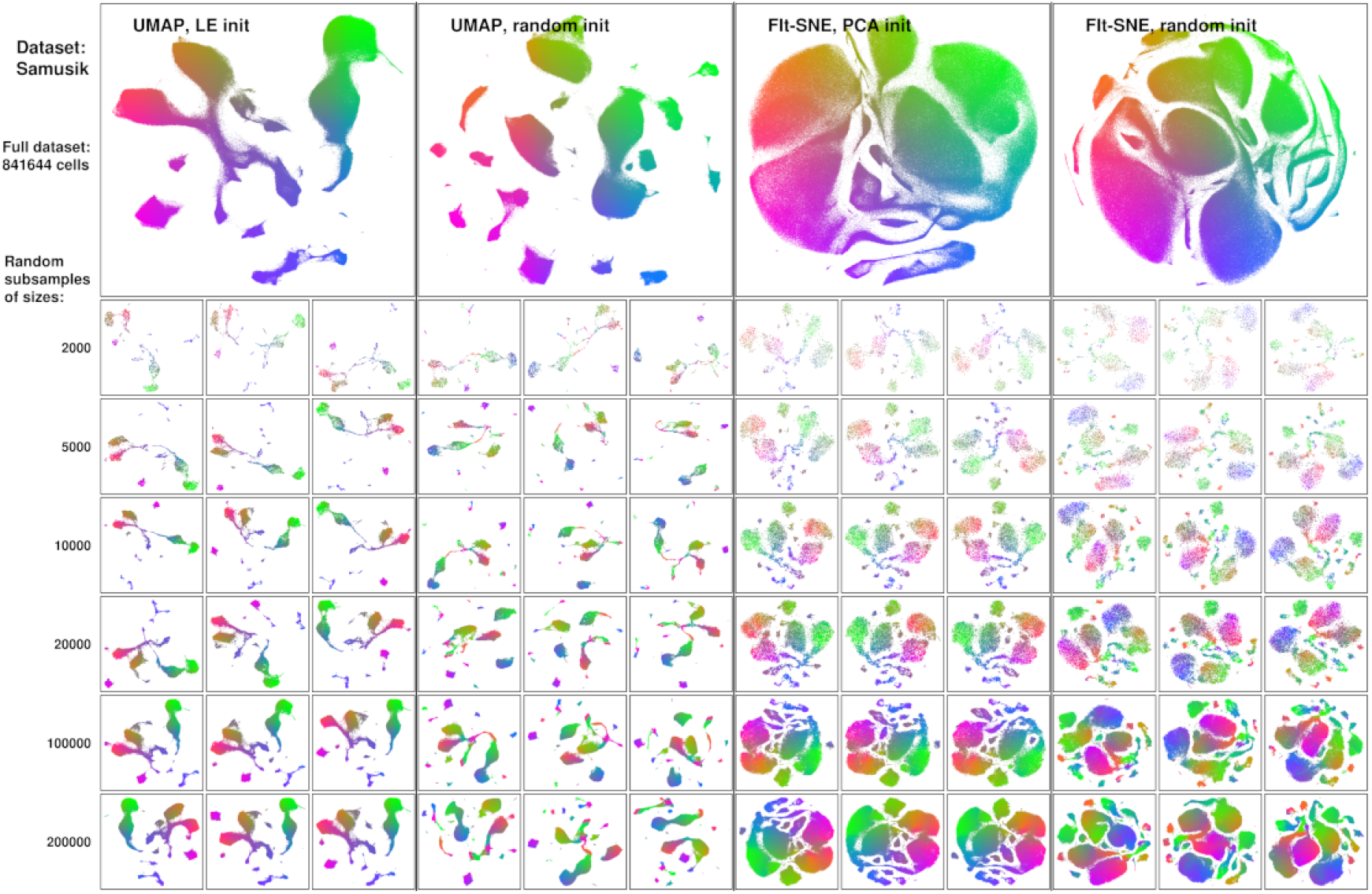
Qualitative assessment of the reproducibility of embeddings. The exact analogue of Supplementary Figure 7a from the original publication.

**Extended Data Figure 4:**
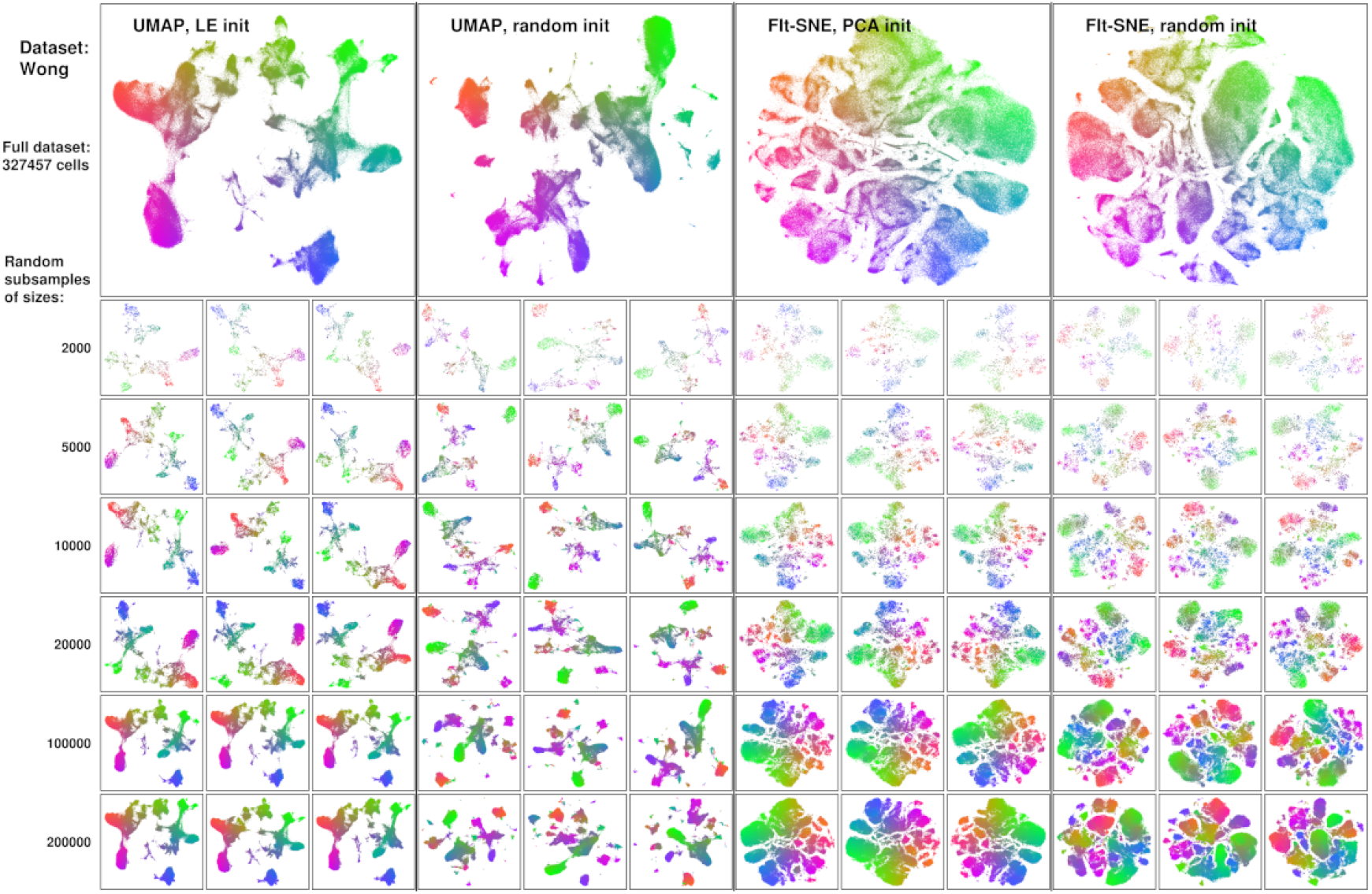
Qualitative assessment of the reproducibility of embeddings. The exact analogue of Supplementary Figure 7b from the original publication.

**Extended Data Figure 5:**
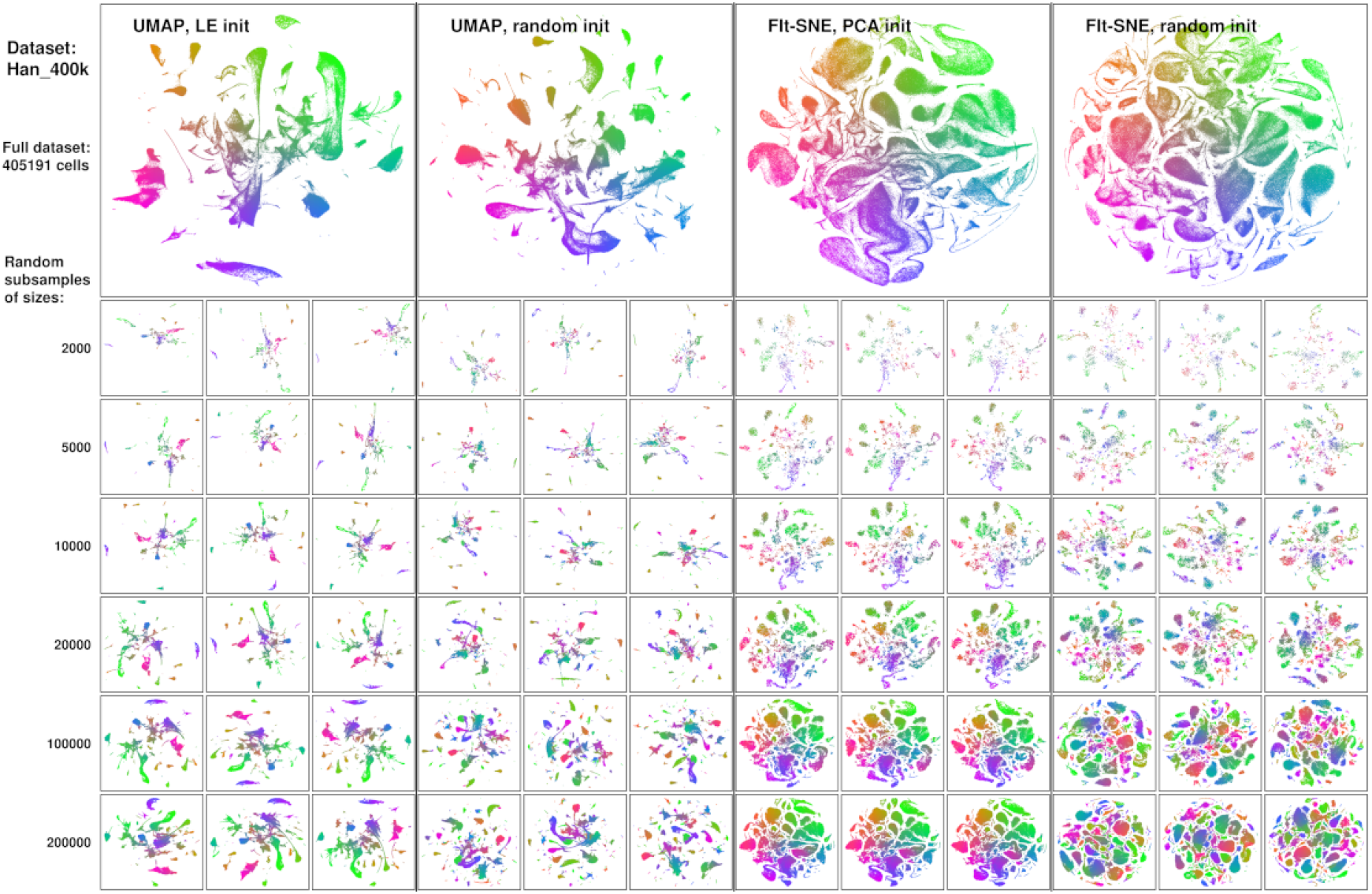
Qualitative assessment of the reproducibility of embeddings. The exact analogue of Supplementary Figure 7c from the original publication.

**Extended Data Figure 6:**
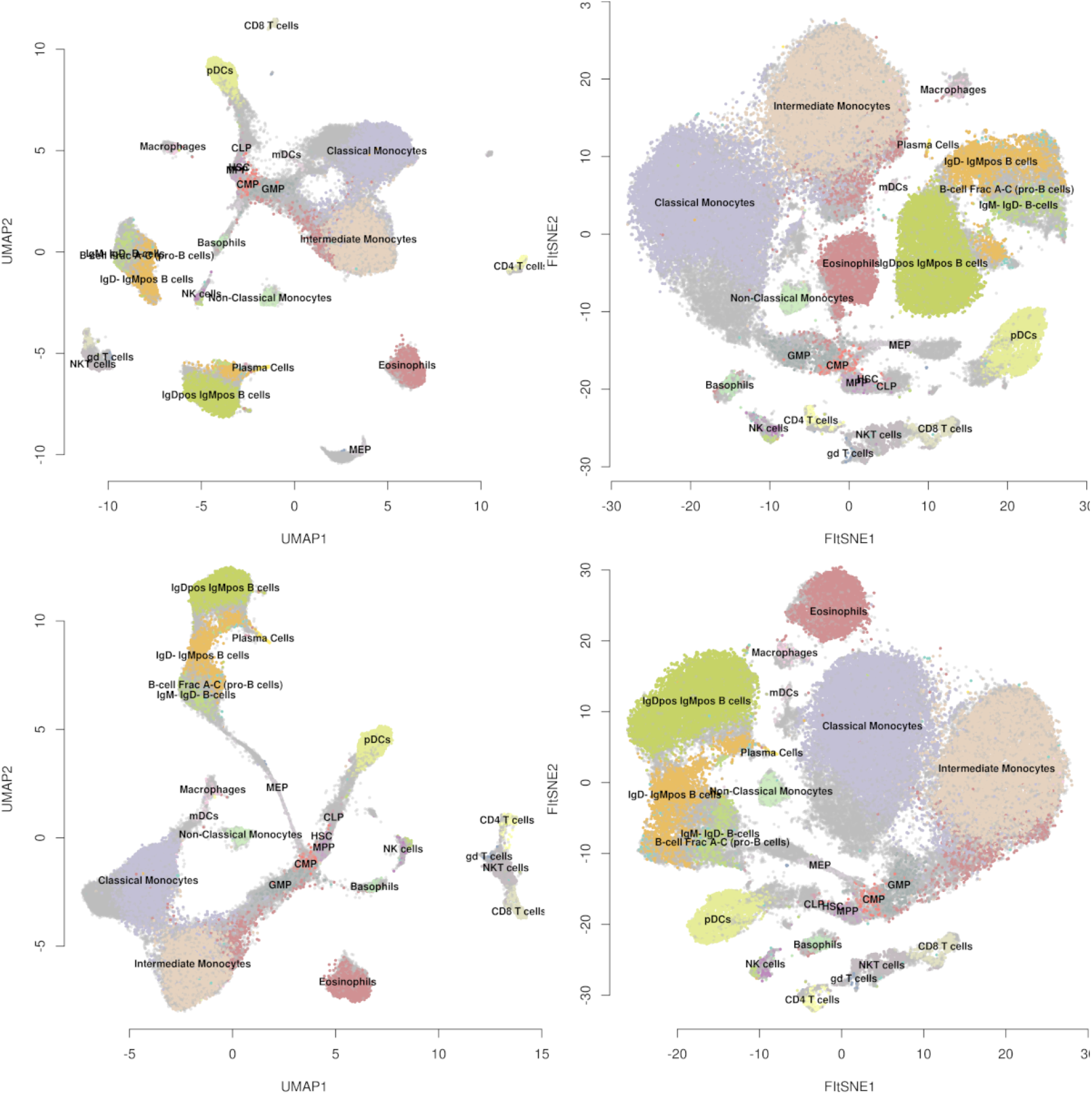
Annotated embeddings of the Samusik_01 dataset (sample size *n*= 86 864). Top row: UMAP with random initialization (left) and t-SNE with random initialization (right). Bottom row: UMAP with default initialization (left) and t-SNE with PCA initialization (right). The bottom-left and upper-right panels are analogues of Figure 2a,b from the original publication. While Becht et al. pointed to t-SNE’s failure to colocalize the T cells in the embedding, we note that with random initialization, UMAP does the same.

